# Spatial mnemonic encoding: Theta power decreases co-occur with medial temporal lobe BOLD increases during the usage of the Method of Loci

**DOI:** 10.1101/052720

**Authors:** M.-C. Fellner, G. Volberg, M. Wimber, M. Goldhacker, M. W. Greenlee, S. Hanslmayr

## Abstract

The Method of Loci is one, if not the most, efficient mnemonic encoding strategy. This strategy combines the core cognitive processes commonly linked to medial temporal lobe (MTL) activity: spatial processing and associative memory processes. During such tasks fMRI studies consistently demonstrate increases in medial temporal lobe (MTL) activity, while electrophysiological studies have emphasized the important role of theta oscillations (3-8 Hz) in the MTL. How MTL activity is linked to theta oscillatory EEG power, however, is unknown. Specifically, it is not known whether increases or decreases in MTL theta power are associated with increased BOLD signal. To investigate this question we recorded EEG and fMRI while participants used the spatial Method of Loci contrasted to the Pegword Method, a similarly associative but non spatial mnemonic. Surprisingly, the more effective spatial mnemonic induced a pronounced theta power decrease in the left MTL compared to the non-spatial associative mnemonic strategy. This effect was mirrored by BOLD signal increases in the MTL. This pattern of results suggests that theta oscillations in the MTL are negatively related to BOLD signal increases. These findings extend the well-known negative relation of alpha/beta oscillations and BOLD signals in the cortex to theta oscillations in the MTL. The results also demonstrate that decreases in theta power can index MTL involvement during encoding.

**Significance Statement:** Studies investigating the oscillatory correlates of memory encoding largely focus on activity in the theta frequency and often implicitly assume that increases in theta activity reflect similar processes as typically reported increases MTL activity changes in fMRI studies. The presented study found decreases in theta power, not increases, closely mapping to MTL BOLD signal increases during the same paradigm. This finding is in line with studies showing a negative relationship between low frequency power and BOLD changes in the cortex, but challenges the assumption that theta power increases reflect MTL activity. The reported findings importantly contribute to answer the question of how and which oscillatory activity indexes MTL memory processes.

## Introduction

Converging work in animals and humans has linked two important cognitive functions to medial temporal lobe structures: spatial processing and memory (Burgess et al., 2002; Buzsaki and Moser, 2013). The influence of spatial processing on memory formation is documented since ancient Greek times: The Method of Loci, a mnemonic strategy based on linking to be learned material to waypoints on a familiar route is an outstandingly efficient strategy to memorize new information (Roediger, 1980). This spatial mnemonic is up to today the preferred strategy of memory athletes memorizing impressive amounts of arbitrary information by associating it to spatial cues (Maguire et al., 2003).

Especially in animals, but also in humans, MTL theta oscillations have been implicated in spatial processing and navigation (Vanderwolf, 1969; Ekstrom et al., 2005; Watrous et al., 2013b). However, concerning memory formation, there is still an ongoing debate about the functional relationship of theta oscillations and MTL activity during memory encoding: are increases or decreases in theta power related to memory formation and MTL engagement? (for a review see, (Hanslmayr and Staudigl, 2014). Several studies report increases in theta power during memory formation during subsequently remembered items in contrast to subsequently forgotten items (Subsequent memory effect, SME) and hypothesized that these increases in theta power reflect MTL involvement (Klimesch et al., 1996; Nyhus and Curran, 2010; Staudigl and Hanslmayr, 2013; Backus et al., 2016). In contrast, other studies found decreased theta activity to be related to successful memory encoding (Burke et al., 2013; Long et al., 2014; Greenberg et al., 2015). Concerning fMRI a more consistent picture emerges where MTL structures are reliably more active during memory tasks (Kim, 2011), especially during tasks that combine associative and spatial processing (Uncapher et al., 2006; Bird and Burgess, 2008; Staresina and Davachi, 2009).

A ubiquitous finding in the cortex is a negative relation between BOLD signal and low frequency power (5 – 30 Hz) (Mukamel et al., 2005; Hanslmayr et al., 2011; Scheeringa et al., 2011; Hermes et al., 2014; Zumer et al., 2014). However, studies investigating theta power changes during spatial processing and associative memory formation suggest a positive relation between MTL activity and theta power (Kaplan et al., 2012; Staudigl and Hanslmayr, 2013; Backus et al., 2016). It therefore remains an open question whether indeed increases or decreases in theta power reflect increases in MTL activity (Lisman and Jensen, 2013).

In order to investigate the relationship between MTL BOLD signal and theta power dynamics we instructed participants to use associative mnemonic encoding strategies: the spatial Method of Loci and the non-spatial Pegword Method (see Figure 1). Both mnemonics entail linking to-be-learned items to internal cues, waypoints in the case of the Method of Loci, and number related pegs in the case of the Pegword Method (Figure 1 A&B). The non-spatial mnemonic was specifically chosen to control for the potential qualitative change in SMEs and MTL involvement during spatial processing. EEG and fMRI was measured in separate groups of participants (Mukamel et al., 2005), because movement-artifacts severely limit the interpretability of theta activity in simultaneous EEG-fMRI (Fellner et al., 2016).

**Figure 1:**
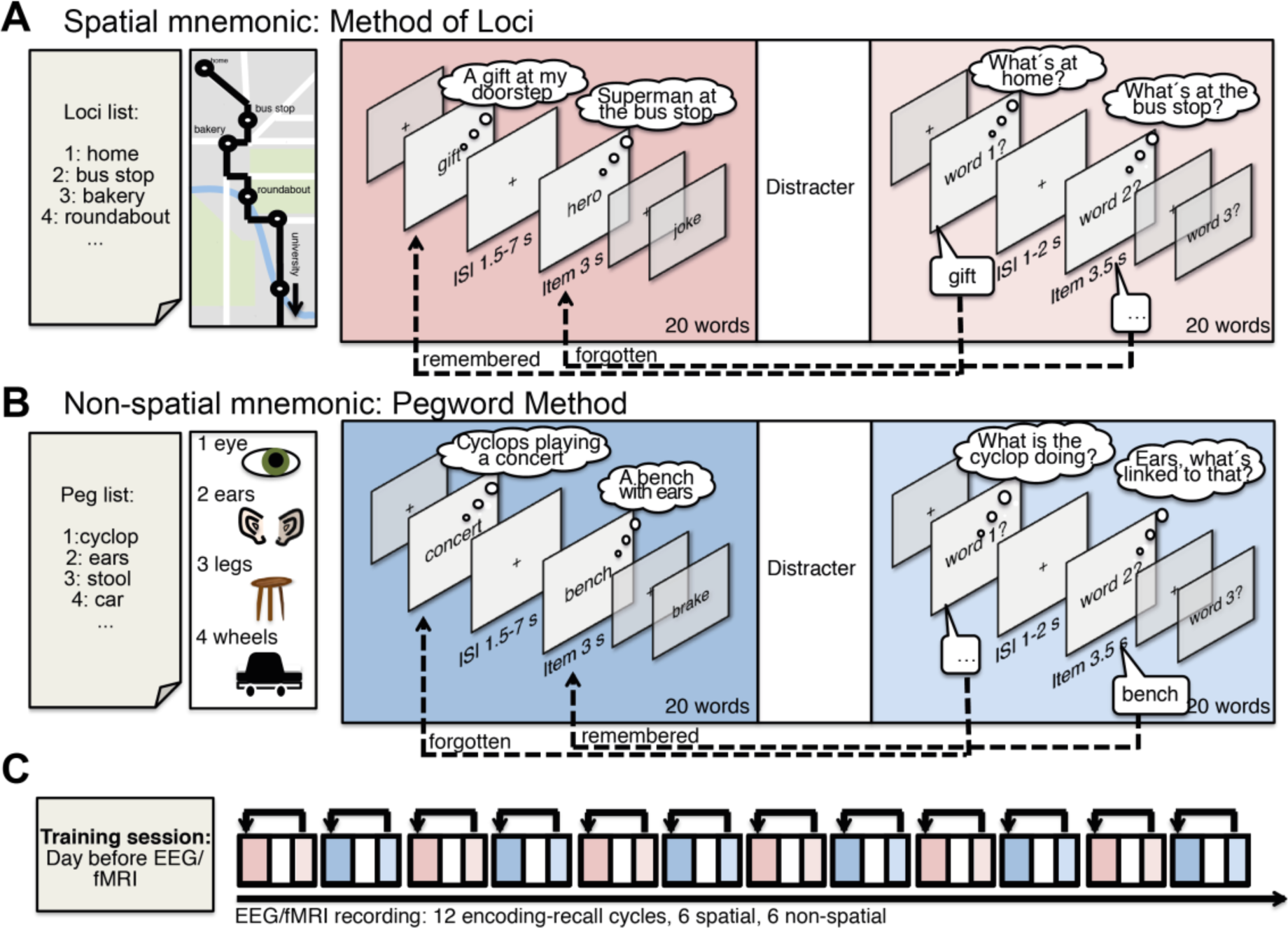
Memory encoding paradigm. Participants were trained to use two mnemonic encoding strategies: the spatial Method of Loci (A) and the non-spatial pegword method (B). In both methods participants have to link internal cues, which are either familiar way points or semantic associations to numbers, to items presented during the encoding phase. During each encoding phase, lists of 20 words were presented sequentially followed by a distracter task and a free recall phase. The whole experiment (C) entailed a training phase the day before and 12 encoding-recall cycles during EEG or fMRI recordings

## Materials and Methods

### Subjects and recording sessions

Two separate groups of participants underwent EEG recording (30 participants) and fMRI scanning (25 participants). Nine EEG datasets had to be excluded (trial numbers below 15 after artifact correction) resulting in a sample of 21 datasets (age 18-24. M=20.19, 12 female). Two fMRI datasets had to be excluded (because of a missing structural scan and task performance two standard deviations below the average) resulting in a sample of 23 datasets (age 18-36, M=22.9, 15 female).

All subjects spoke German as their native language, reported no history of neurologic or psychiatric disease, and had normal or corrected to normal vision. All participants gave their written informed consent, and the experimental protocol was approved by the local ethical review board. During fMRI scanning, simultaneous EEG was recorded. Major MR scanner induced EEG artifacts, especially in the theta frequency range, prevented reliable analysis of the simultaneously recorded EEG (see (Fellner et al., 2016) for a discussion of these artifacts).

### Task design

The exact same paradigm was presented to participants during fMRI and EEG. In the encoding phase 20 to-be-learned words were presented sequentially (word presented for 3sec followed by fixation cross shown for 1.5-7 sec exponentially jittered). Participants were instructed to associate each word with the corresponding cue in the loci or pegword sequence. Using the spatial Method of Loci they were instructed to visualize to-be-learned items (e.g. first item gift) at the respective waypoint (e.g. first loci cue home). For instance participants might imagine a nicely wrapped gift at their doorstep (Figure 1A). During the non-spatial pegword method, they were asked to focus on semantic relations between the item (e.g. first item concert) and the respective pegword (e.g. first pegword cyclop), such that a possible association could be “A band of cyclops playing a concert” (Figure 1B). During recall, participants were instructed to use the pegwords and loci way points as retrieval cues to recall the words in the same sequence as during the encoding phase. Participants were asked to recall the 20 words of the prior encoding phase in their original order, whenever the screen showed a “word no. x” cue. Only words recalled in the corrected order were included as remembered trials, words not recalled were labeled forgotten.

The fMRI-or EEG-recorded part of the experiment consisted of 12 repeated memory encoding and recall cycles, with each encoding-recall block consisting of an encoding phase, a visual detection task, a free recall memory test and a short 20 sec rest period (see Figure 1C). Each experiment was split into 4 consecutive recording sessions to keep file sizes manageable; the MR image acquisition was thus stopped after every third encoding-recall cycle. A visual detection task (ca. 2.5 min) was serving as a distracter task in order to (similar task as in (Hanslmayr et al., 2013)). In-scanner verbal responses were recorded using an fMRI-compatible microphone (MRconfon). Scanner noise was removed from the resulting audio files using the free software package Audacity (http://audacity.sourceforge.net/). For two participants, data from one of the 4 fMRI sessions had to be discarded because of faulty microphone recordings. During the EEG experiment, recall performance was scored manually by the experimenter.

A training session took place one day before fMRI or EEG recording to ensure proper usage of both mnemonics. Prior to training, participants received basic information about mnemonics and were instructed to prepare 20 waypoints on a highly familiar path (e.g. the way to the university), and to memorize the 20 pegword cues. The training session then consisted of 3 parts: First, memorization of all loci and pegword cues was tested. Second, participants had to encode an exemplary list of 20 words naming all employed associations between words and the loci or pegword cues. If the chosen associations did not fit with the task instruction, the training session was restarted. Third, 4 practice encoding-recall-cycles with same timing as during EEG and fMRI were carried out.

As study material, 360 words were drawn from the MRC Psycholinguistic Database (Coltheart, 2007), translated into German and separated into 16 lists with 20 words each. Four lists were used during the training session; the other 12 lists were used during the EEG and fMRI recordings. Each of these 12 lists was matched according to average word frequency (M=61.98; SEM=1.58), number of letters (M=5.59; SEM=0.04), syllables (M1.84; SEM=0.02), concreteness (M=375.65; SEM=2.17), and imageability (M=414.21; SEM=2.83). Word lists were counterbalanced across participants and encoding tasks. Word order in each list was randomized.

### EEG recording

The EEG was recorded from 63 channels in an equidistant montage, (EasyCap, Herrsching, Germany, BrainampMR). Recordings were referenced to Fz and later re-referenced to average reference. Impedances were kept below 20 kΩ. The signals were amplified between 0.1–250 Hz. The EEG data were sampled at 500 Hz with an amplitude resolution of 0.5μV

### EEG preprocessing and analysis

All EEG data analyses were carried out using custom MATLAB scripts and fieldtrip (http://www.fieldtriptoolbox.org, (Oostenveld et al., 2011)). Data was epoched in trials −2.5 to 3.5 sec around each item onset during encoding. Data was visually inspected to exclude trials with idiographic artifacts (channel jumps, muscle artifacts, noisy channels) from further analysis. Noisy channels were excluded (in 4 datasets up to 3 electrodes were excluded). Infomax independent component analysis was applied to correct for residual artifacts (e.g. eye blinks, eye movements, or tonic muscle activity). On average 3.8 ICs were discarded (range: 1-8 ICs). Data of rejected channels were interpolated using neighboring electrodes. On average 62.9 spatial/remembered trials (range 33-96), 45.7 non-spatial/remembered trials (range 23-74), 34.7 spatial/forgotten trials (range 19-61), and 49.7 non-spatial/forgotten trials (range 22-81) passed artifact corrections.

Data was filtered using wavelets with a 5 cycle length to obtain oscillatory power between 2 Hz and 30 Hz. Resulting data was z-transformed to respective mean and standard deviation of power across the time dimension i.e. across all trials of each frequency band and channel. For each subject and condition (remembered, forgotten, loci, pegword) all trials were averaged and smoothed with Gaussian kernel (FWHM 200ms and 2 Hz) to attenuate inter-individual differences and to control for the time-frequency resolution trade-off across frequencies.

Source analysis was carried out using a linearly constrained minimal variance (LCMV) beamformer (Van Veen et al., 1997), calculating a spatial filter based on the whole length of all trials. For all subjects, a standard source model with a grid resolution of 12 mm based on the Montreal Neurological Institute (MNI) brain and standard electrode positions realigned to the MNI MRI was used. The source time-course for each grid point was calculated and subjected to a wavelet analysis and z transformed as for electrode level data. For virtual electrode analysis, data across all grid voxel covering the ROI were averaged. Grid-voxel data was interpolated to a 2mm resolution single subject MNI brain for plotting and to define locations of clusters and peaks.

For statistical analysis, non-parametric cluster permutatieon tests were employed as implemented in fieldtrip (Maris and Oostenveld, 2007). First, clusters of coherent t-values exceeding a certain threshold (here p<0.05) along selected dimensions (time, frequency, electrodes/grid voxels) are detected in the data. Summed t-values of these data clusters are then compared to a null distribution of t-sums of randomly obtained clusters by permuting condition labels across subjects. This procedure effectively controls for type-I-errors due to multiple testing. Clustering was employed along different dimensions depending on whether data was averaged based on prior windows of interest: for testing scalp level effects in the theta band range three-dimensional clustering was employed (time x frequency x electrodes), analysis in virtual electrodes employed a two-dimensional clustering (time x frequency). Analysis of whole brain source localized effects was carried out for the average activity in the time-frequency windows identified in scalp analysis clustering along the spatial dimension.

For identification of time-frequency windows showing significant differences between conditions across all lower frequencies (1-30 Hz) in a more explorative/liberal manner a sliding window cluster permutation test was used (for details (Staudigl and Hanslmayr, 2013)). A cluster permutation test here is calculated for each 300 ms x 1 Hz time-frequency bin, clustering across electrodes. This approach yields a p-value for each of these time-frequency bins. For time-frequency windows showing coherent bins of significant effects, an additional cluster permutation test was carried out to identify spatial topographies of these effects.

### fMRI recording

Imaging was performed using a 3 Tesla MR head scanner (Siemens Allegra). During fMRI scanning, 2475-2480 whole-brain images, consisting of 34 axial slices, were continuously acquired using an interleaved, standard T2*-weighted echo-planar imaging sequence (repetition time TR=2000 ms; echo time TE=30 ms; flip angle=90°; 64×64 matrices; in-plane resolution: 3×3 mm; slice thickness: 3 mm). High-resolution (1mm isotropic voxel size) sagittal T1-weighted images were acquired after the functional scans, using a magnetization-prepared rapid gradient echo sequence (TR=2250 ms; TE=2.6 ms) to obtain a 3D structural scan.

### fMRI preprocessing and analysis

Image preprocessing and statistical analysis was performed using SPM8 (Wellcome Department of Cognitive Neurology, London: UK, www.fil.ion.ucl.ac.uk/spm), running on MATLAB (Version 2012b, The MathWorks, Natick, MA). After discarding the first two images of each session, time series were corrected for differences in slice acquisition time, spatially realigned to the first image of the session, and unwarped. The mean functional image was co-registered to the structural image. Global effects in the functional time series within each session and voxel were removed using linear detrending (Macey et al. 2004) All functional images were then normalized to MNI space (Montreal Neurological Institute, www.mni.mcgill.ca) using the normalization parameters determined from segmentation of the structural image. As a last step images were smoothed with a Gaussian kernel of 8 mm (FWHM).

Scans of all four session were concatenated in first level GLMs. Activity related to subsequent memory and encoding task was modeled by event related stick regressors for each condition (loci-remembered, loci-forgotten, pegword-remembered, pegword-forgotten) convolved with the canonical first order hemodynamic response function. Further regressors of no interest were modeling the rest periods, free recall periods (separately for each encoding condition), and the distracter task. Session-specific regressors, linear drifts within each session, and movement parameters determined during realignment were also included in the model. Contrasts capturing encoding effects for the 4 conditions of interest were calculated in each single participant, and combined in a 2 (task) × 2 (subsequent memory) full factorial random effects model on a group level.

As a first step, a ROI analysis on MTL effects was carried out using small volume correction on bilateral MTL defined by Wake Forest University WFU pick atlas (https://www.nitrc.org/projects/wfu_pickatlas/, parahippocampal cortices plus hippocampi using TD atlas, the same ROI definition as used for virtual electrodes ROI analysis). Small volume corrected effects are reported using a p<0.001, cluster size>10 voxels. An additional whole-brain analysis was carried out using a threshold of p<0.001 uncorrected, cluster p-level, family-wise error corrected <0.05. Marsbar (http://marsbar.sourceforge.net) was used to extract parameter estimates of significant clusters. Results are plotted on the mean normalized structural scan of all subjects.

## Results

### Behavioral performance

Memory performance in EEG and fMRI was reasonably high, in line with the efficiency of mnemonic strategies (Roediger, 1980). Memory performance (Figure 2) in both datasets was significantly higher during the spatial encoding than during the non-spatial encoding task (fMRI: t(22)=6.26, p<0.0001, EEG: t(20)=10.23, p<0.0001). This was a very robust effect, which was visible in almost every single subject: 19 of 21 EEG participants and 20 of the 23 fMRI participants showed higher recall rates using the spatial mnemonic, demonstrating the power of the Method of Loci as a mnemonic (Roediger, 1980).

**Figure 2:**
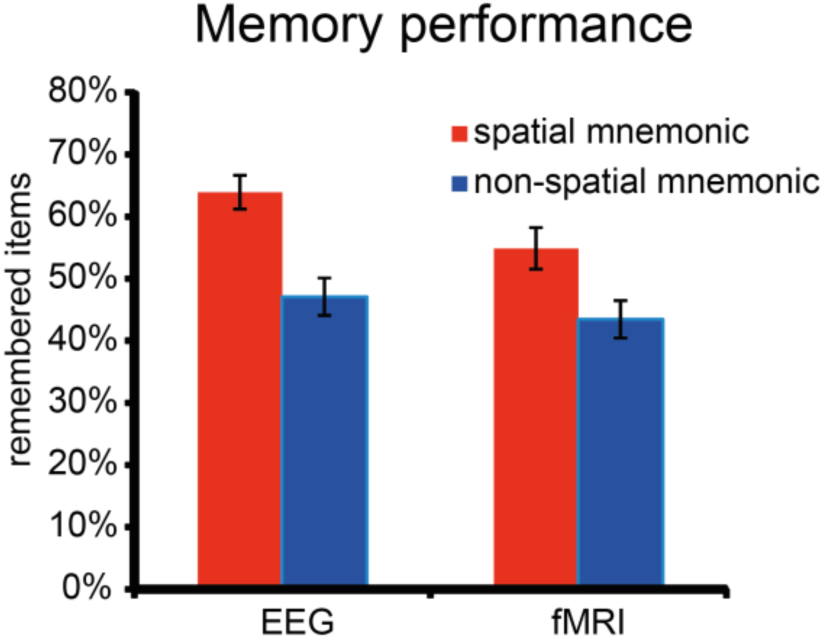
Memory performance. Percent of recalled words in the spatial and non-spatial encoding condition separately for the EEG experiment and fMRI experiment. In both datasets memory performance was higher using the spatial Method of Loci mnemonic. Error bars show standard error of the mean.

### EEG scalp level

The first step of analysis focused on theta frequency range as this frequency band has been previously linked to spatial processing and memory formation. Therefore we contrasted theta power during the spatial and non-spatial mnemonic and for remembered and forgotten trials (SME, Figure 3A). A robust theta power decrease for the spatial mnemonic contrasted to the non-spatial encoding strategy was found spanning the whole trial epoch (−1.5 to 2.5 sec, p_corr_ =0.002, Figure 3A). This theta power decrease was ongoing throughout the trial, and does not appear to be triggered by the onset of the to-be-encoded words. Decreases in theta power were also related to successful memory formation as revealed by negative SMEs. A significant cluster of memory related theta decreases was evident from stimulus onset lasting until the end of the trial (0-2.5 sec, p_corr_=0.014, Figure 3A). The extent of the clusters is shown in Figure 3A (left for spatial vs. non-spatial; right for SMEs). This pattern of results shows that spatial mnemonic processing as well as successful memory formation are reflected in theta power decreases.

**Figure 3:**
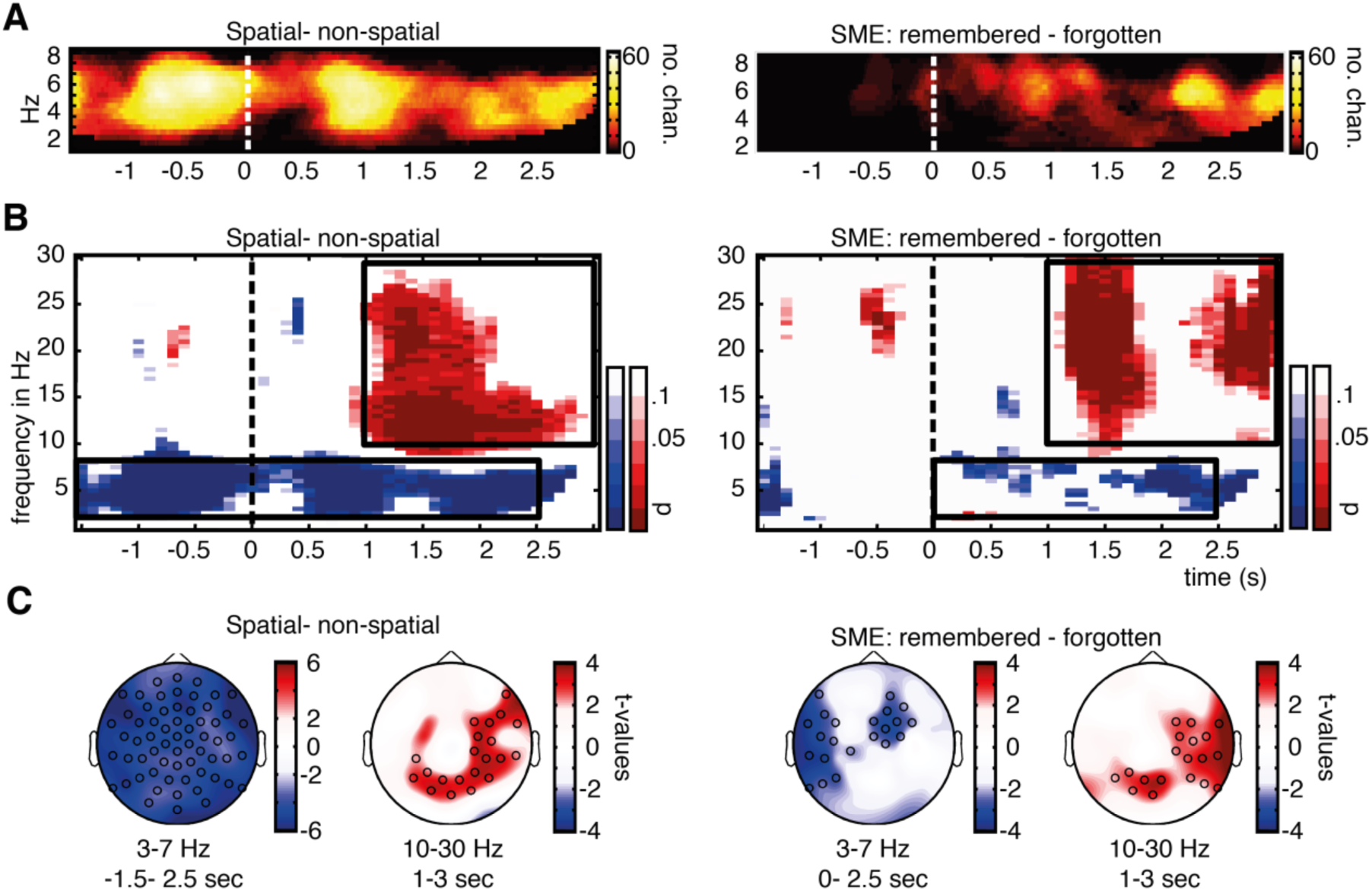
EEG sensor level results. (A) A cluster permutation statistic restricted to the theta frequency range revealed ongoing decreases in theta oscillatory power for spatial mnemonic processing in contrast to non-spatial processing, and item related theta power decreases correlating with successful memory formation. The time-frequency plots show the number of electrodes belonging to the cluster at a certain time-frequency bin in order to visualize the extent of the three-dimensional cluster in the time-frequency dimensions. (B) Additional increases in alpha/beta power during spatial encoding and memory formation were evident after word presentation. Time frequency plots here show p-values of separately calculated cluster permutation tests of each time-frequency bin. (C) Topographies of theta and alpha/beta power effects for the time-frequency windows in which effects were present highlighted in (B, dashed boxes) are plotted below, circles highlight electrodes belonging to a significant cluster (C). Warm colors in (B) & (C) indicate increases in power for spatial processing and successfully encoded items, cold colors indicate decreases in power for spatial processing and successfully encoded items in contrast to non-spatial processing and subsequently forgotten items, respectively.

An additional sliding cluster permutation statistic was carried out on all lower frequency bands (1-30 Hz, Figure 3B) and revealed also effects outside theta range. Significant effects were also obtained in the alpha/beta range (10-30 Hz Figure 3B). Here, a stronger power increase was evident after stimulus presentation (1-3 sec) for the spatial mnemonic relative to the non-spatial strategy. Similarly a positive alpha/beta SME indicating higher alpha/beta power during successful memory formation was evident (10-30 Hz, 1-3sec Figure 3B).

For a first identification of potential sources of EEG effects, topographies of the time windows of significant effects are shown in Figure 3C. The decrease in theta power related to the spatial mnemonic showed a widespread topography with the strongest effects over lateral electrodes (p_corr_=0.001). Theta power decreases related to memory encoding were evident in left temporal lateral and right frontal regions (two separate clusters: p_corr_=0.005, p_corr_=0.011, respectively). Increases in alpha/beta power stronger during the spatial mnemonic were found over parietal electrodes (p_corr_=0.003). Memory related alpha/beta power increases showed a similar posterior and right lateralized topography (two clusters: p_corr_=0.011, p_corr_=0.035).

To investigate potential differences of encoding effects between the spatial and non-spatial mnemonic strategies, interaction effects were calculated by contrasting SMEs of both conditions. No significant clusters were found in a cluster analysis in the theta frequency range (2-10 Hz, similar analysis as for Figure 3A), an additional sliding cluster statistic including all lower frequencies (2-30 Hz similar analysis as for Figure 3B) revealed no coherent effects exceeding a size of 8 time-frequency bins, suggesting that successful encoding with both mnemonic strategies similarly relies on theta power decreases and that indeed theta decreases are related to associative memory encoding and are no a specific correlate of the Method of Loci mnemonic.

### fMRI results

As a first step we investigated MTL BOLD. To this end, a region of interest (ROI) analysis on anatomically predefined MTL regions was carried out (Figure 4A, left & Table 1). BOLD activity increased significantly in bilateral parahippocampal regions during the spatial mnemonic in contrast to the non-spatial task. Positive SMEs were also found in the left MTL (Figure 4A, right & Table 1). In line with our hypotheses, these results suggest that the spatial mnemonic and successful associative memory formation both rely on increased hemodynamic MTL activity.

**Figure 4:**
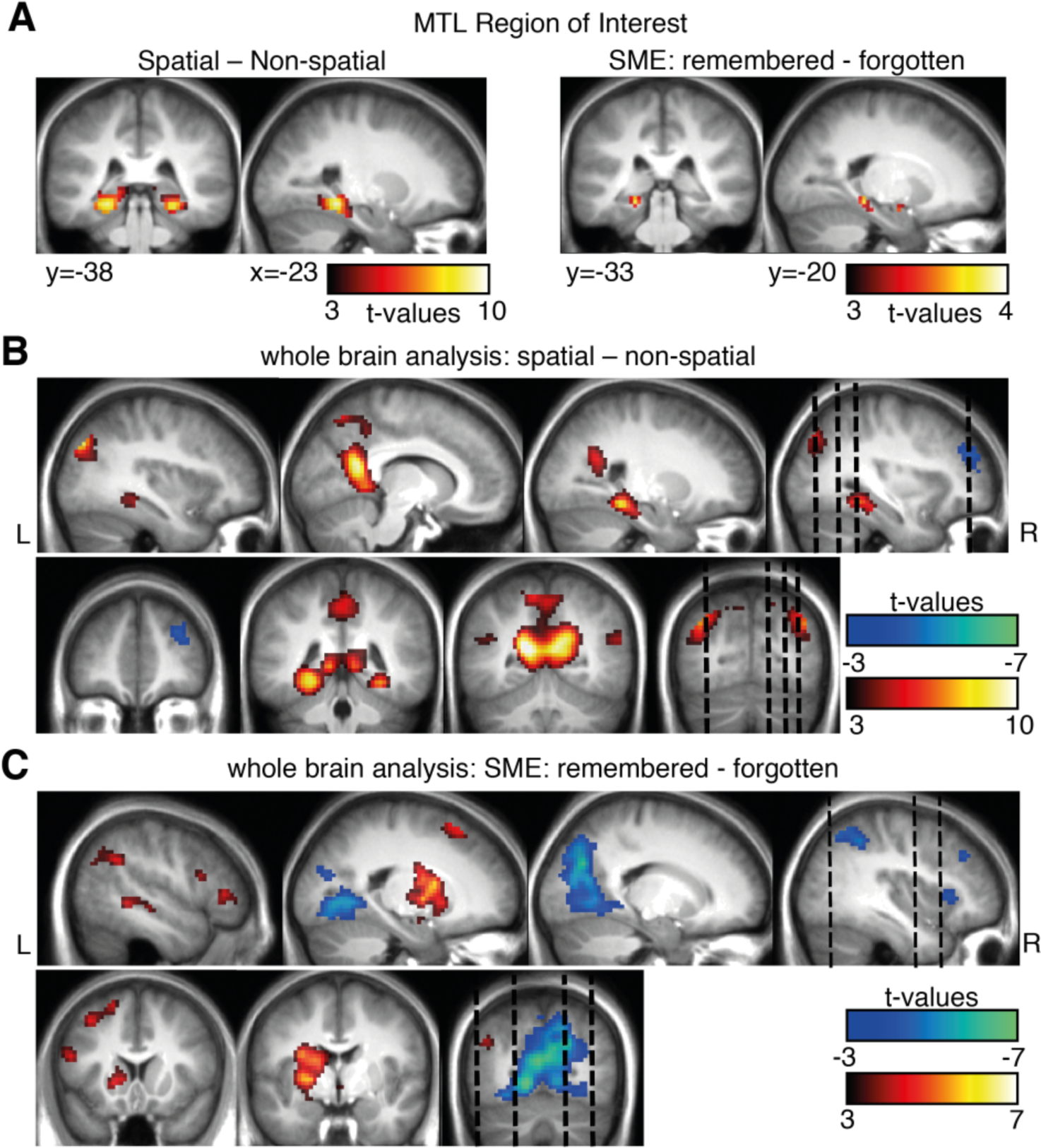
fMRI results for spatial vs. non-spatial contrasts and memory effects. (A) A region of interest analysis was carried out for MTL regions revealing increases in activity for the spatial mnemonic and successful memory formation ( p<0.001, cluster size>10). An exploratory whole brain analysis revealed additional effects in typical spatial cortical networks (i.e. retrosplenial cortex, bilateral MTL, B) and memory related regions (i.e. left inferior frontal gyrus C) (p<0.001, all p<0.05 FWE cluster level). Warm colors indicate higher BOLD signal for spatial processing and later remembered items, cold color indicate higher BOLD signals for non-spatial processing and subsequently forgotten items.

**Table 1:** Locations of peak activation revealed in MTL ROI analysis

| Spatial > Non-spatial |  |  |  |  |  |  |  |
| --- | --- | --- | --- | --- | --- | --- | --- |
|  | hs | BA | size | x | y | z | t |
| Parahippocampal gyrus | L | 36 | 197 | -24 | -40 | -11 | 9.67 |
|  | L |  |  | -9 | -46 | 1 | 7.28 |
| Parahippocampal gyrus | R | 36 | 156 | 27 | -37 | -14 | 8.54 |
| Parahippocampal gyrus | R |  | 20 | 12 | -46 | 1 | 6.83 |
| positive SME: remembered > forgotten |  |  |  |  |  |  |  |
|  | hs | BA | size | x | y | z | t |
| Parahippocampal gyrus | L |  | 14 | -27 | 2 | -14 | 4.07 |
|  | L | 28 |  | -15 | -10 | -17 | 4.00 |
| Parahippocampal gyrus | L | 35 | 18 | -21 | -34 | -11 | 3.79 |
|  | L | 35 |  | -24 | -28 | -20 | 3.38 |

An additional exploratory whole brain analysis (Figure 4B&C & Table 2) was carried out to investigate which other regions are involved during mnemonic encoding. Regions typically associated with spatial processing (Burgess et al., 2002; Epstein, 2008) showed BOLD signal increases during spatial compared to non-spatial processing (Figure 4B): bilateral retrosplenial cortex (posterior cingulate cortex, BA30), and lateral temporal areas of the angular gyrus (BA 39). In contrast the right middle frontal gyrus (Figure 4B) showed relative increases in activity during the non-spatial mnemonic, potentially related to enhanced control processes (Schott et al., 2005). Positive subsequent memory effects (Figure 4C), irrespective of task, were found in the left hemisphere in inferior frontal gyrus, an area typically involved in memory encoding of verbal material (Kim, 2011). Additional memory related activity was evident in striatal areas (putamen, caudate body), left superior and middle temporal gyrus. Relative decreases in BOLD signals during remembered words contrasted to forgotten words, i.e. negative SMEs, were evident in bilateral occipital areas (cuneus, lingual gyrus) and right lateralized parietal and frontal regions (Figure 4C).

**Table 2:**
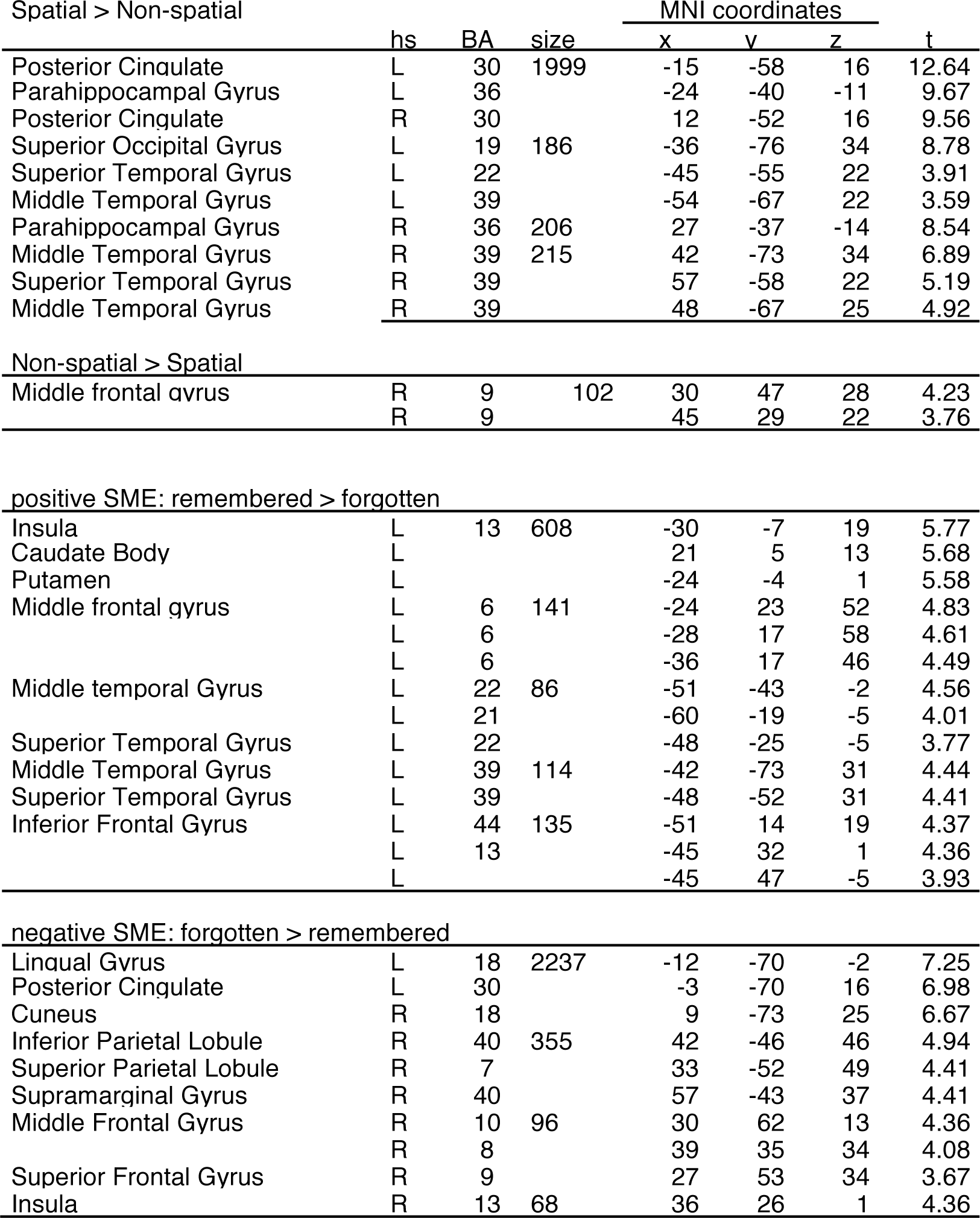
Locations of peak activation revealed in the whole brain analysis

To elucidate if memory encoding related activity differs between the spatial and non-spatial mnemonic strategies, interaction effects were calculated. No significant differences in memory encoding between encoding strategies (i.e. no interaction effects between condition and memory) were found in the ROI analysis or in the whole-brain analysis (p<0.001, cluster size > 10). Spatial and non-spatial encoding therefore seems to rely on BOLD activity in similar brain regions. This absence of differences in encoding related activity in fMRI results closely matches the absence of differences in EEG findings.

### EEG source analysis

Based on our hypotheses and backed up by the fMRI results we placed virtual electrodes in bilateral MTL regions to investigate theta power on source level. Oscillatory source effects were estimated by means of a source localization analysis utilizing LCMV beamforming (Van Veen et al., 1997) and virtual electrode placement was based on the same region of interest as for fMRI ROI in left and right MTL (see Figure 5A). In bilateral MTL theta power showed stronger decreases pre-and post-stimulus during spatial mnemonic processing than during the non-spatial strategy (left MTL, two clusters p_corr_=0.001 & p_corr_=0.001, right MTL two clusters p_corr_=0.001 & p_corr_=0.001, Figure5B). SMEs in the theta band were only evident in the left MTL: theta power decreases were significant post-stimulus (two clusters p_corr_=0.014 & p_corr_=0.022, Figure 5B), but not in the right MTL (all cluster p_corr_>0.15). This pattern of bilateral MTL effects during spatial processing and left lateralized MTL effects during memory formation parallels the fMRI MTL results (see Figure 4A).

**Figure 5:**
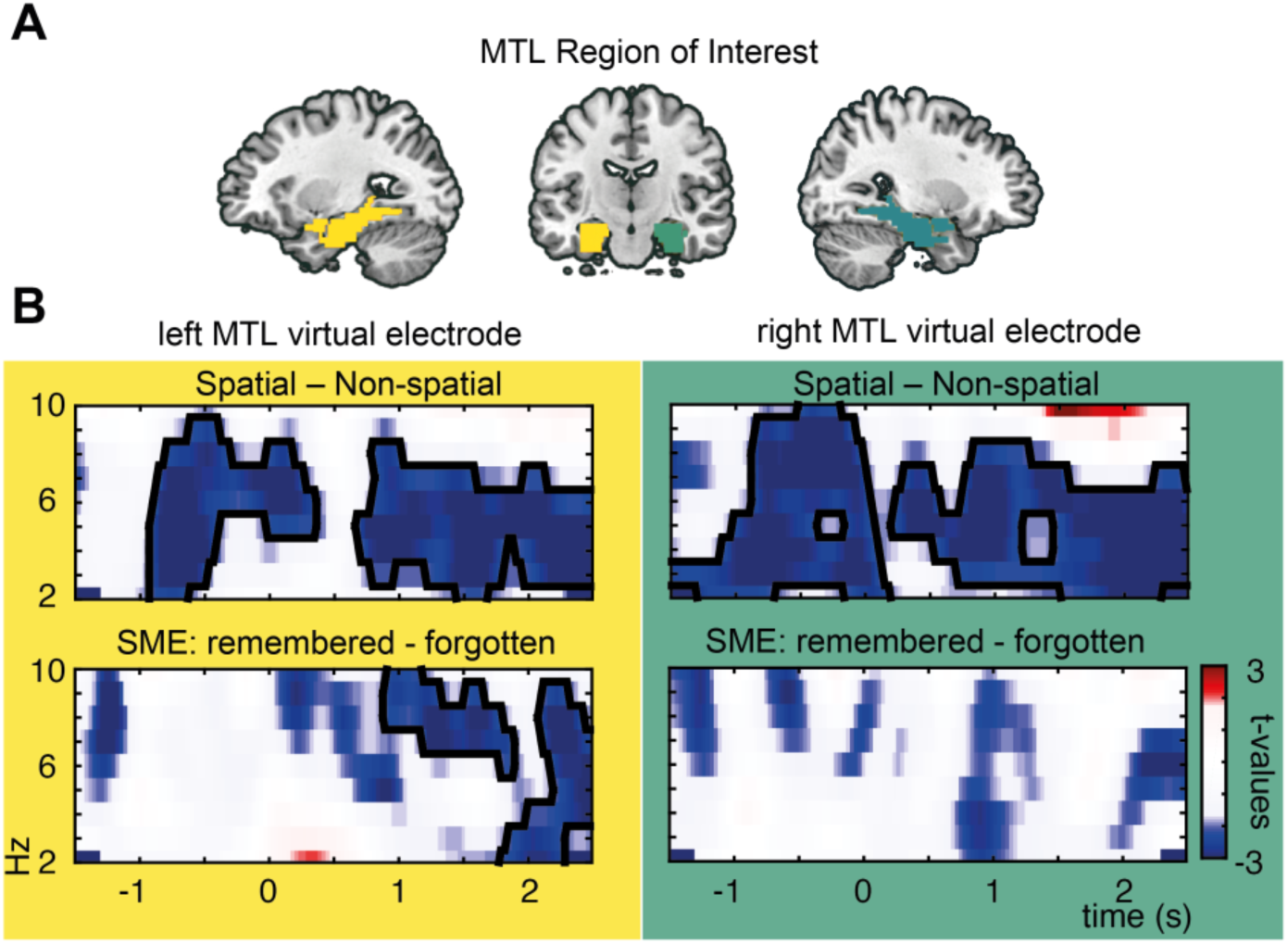
Theta power changes in MTL. The same region of interest here highlighted in green and yellow for virtual electrode EEG activity and used for fMRI analysis. (D) Theta power effects of virtual electrodes in left and right MTL, theta power decreases were found bilaterally for spatial vs non-spatial processing and left lateralized for successful memory formation. Significant clusters are highlighted by black contours.

To illustrate the whole brain distribution of power effects, peaks of source localized task and memory related oscillatory activity are shown in Figure 6. Theta power decreases for spatial mnemonic processing compared to non-spatial processing were strongest in the left anterior MTL (p_corr_=0.001, t_peak_=−7.2, peak MNI coordinate: x=−30, y= −19, z= −32, parahippocampal gyrus; Figure 6A). Memory formation related theta power decreases were strongest in left lateral temporal lobe areas (p_corr_=0.002, t_peak_=−5.9, peak MNI coordinate: x= −56,y= −44, z=−7, middle temporal gyrus, Figure 6A). To further characterize the extent of effects in source space, areas including at least ~1.5 grid voxels (i.e. 324 interpolated voxels) extending a t-threshold set relative to the peak t-value are summed up in table 3. Note the overlap of theta power decreases and BOLD activity increases in MTL and temporal regions (cf. Figure 6A & Figure 4).

**Figure 6:**
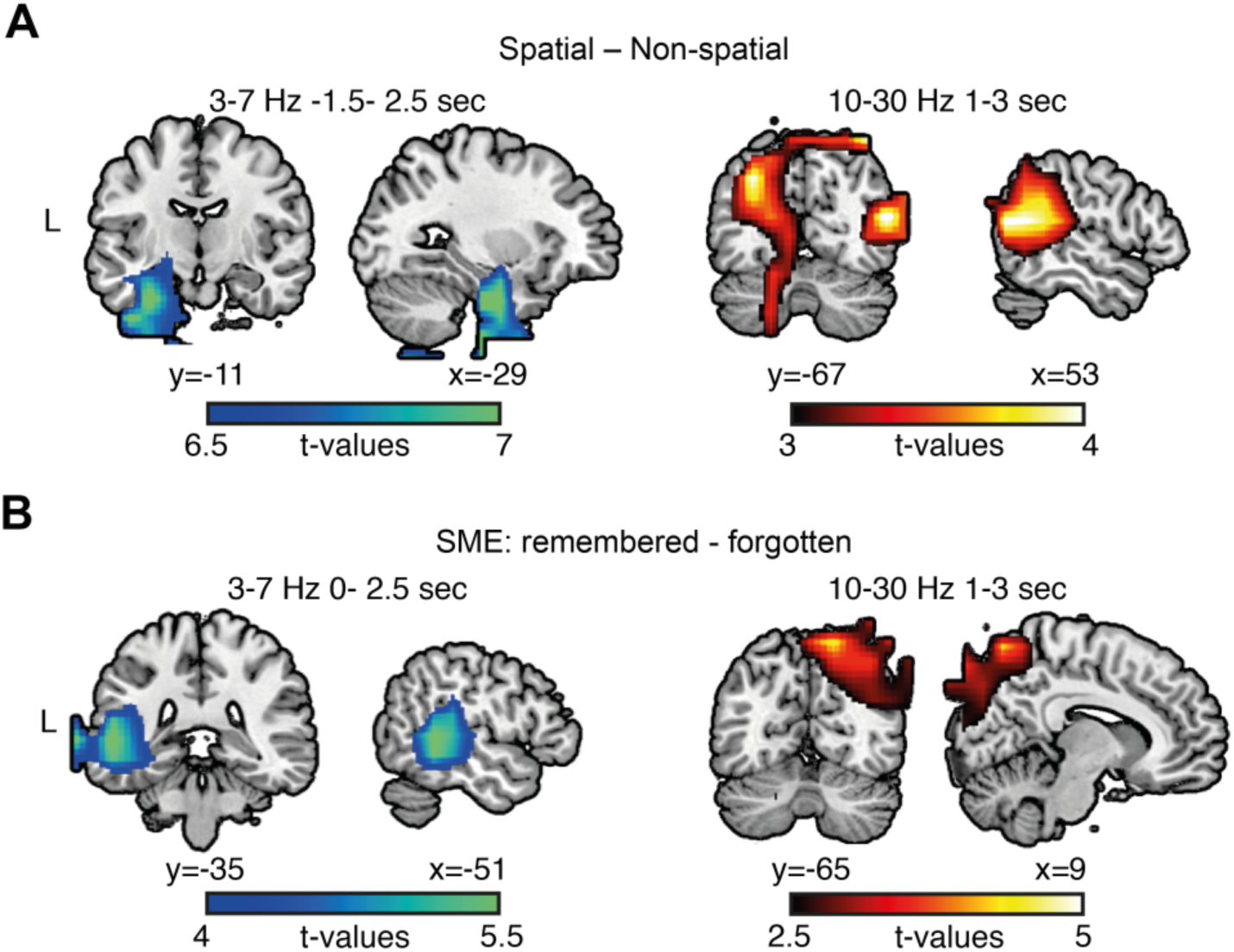
Peaks of EEG source localization results. (A) Decreases in theta power for spatial processing were strongest in anterior MTL areas (peak left parahippocampal gyrus), Alpha/beta power increases were strongest in occipital-parietal-temporal areas for spatial vs. non-spatial processing (peak in posterior middle temporal gyrus)(B) Theta power decreases during successful memory formation were strongest in left temporal areas (peak in middle temporal gyrus). Increases in alpha/beta power during memory formation were found in occipital-parietal areas (peak in superior parietal lobule)

**Table 3:**
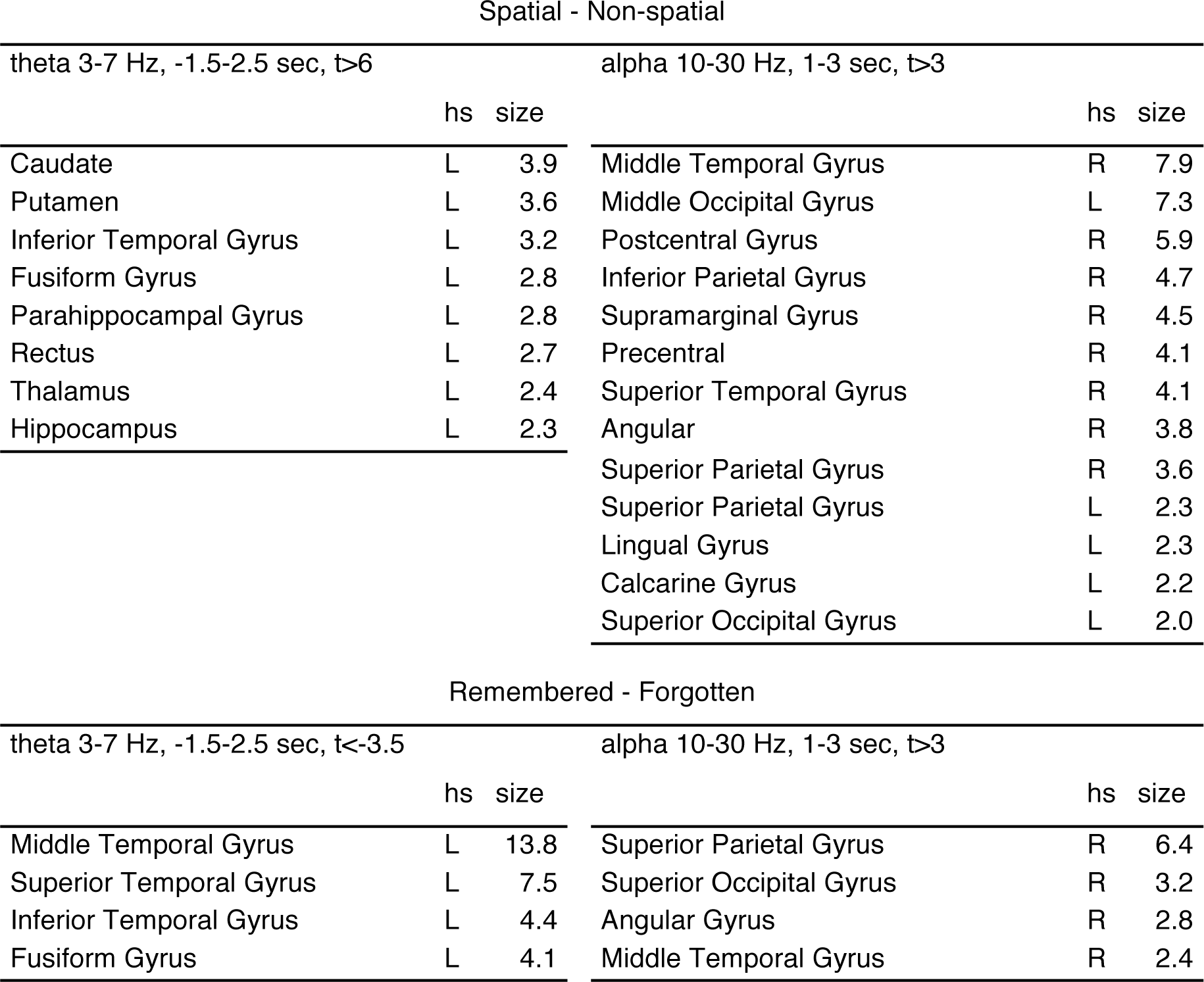
Anatomic labels of EEG source contrasts

Anatomical labels were derived by inte rpolati ng the low spatially resolved source grid res ults (12mm^3^) onto a highly spatially resolved MNI template brain (2 mm^3^). Only areas including more template voxels as in 2 significant grid voxels (i.e. 436 template brain voxels) are reported here, size refers to the number of grid voxels (12mm^3^) exceeding the given t-threshold.

Alpha/beta power increases for spatial mnemonic processing and positive SMEs were found in occipito-parietal areas and right lateralized regions (spatial-non-spatial: p_corr_=0.012, t_peak_=4.3, peak MNI coordinate: x= 52,y= −66, z=17, middle temporal gyrus, SME: p_corr_=0.038, t_peak_=4.7, peak MNI coordinate: x= 18,y= −67, z= 64, superior partial lobule, Figure 6B). Similarly to theta decreases and BOLD increases, alpha/beta power and BOLD activity seem to be negatively related: increases in alpha/beta power overlap with areas were BOLD decreases during successful memory formation were evident (cf. Figure 6B & Figure 4C).

To further illustrate the relationship between the MTL BOLD signal and theta, average EEG theta power and the corresponding fMRI beta estimates derived from the left MTL ROI are plotted in Figure 7. Decreases in theta power mirror fMRI effects such that decreases in theta power are associated with MTL BOLD signal increases.

**Figure 7:**
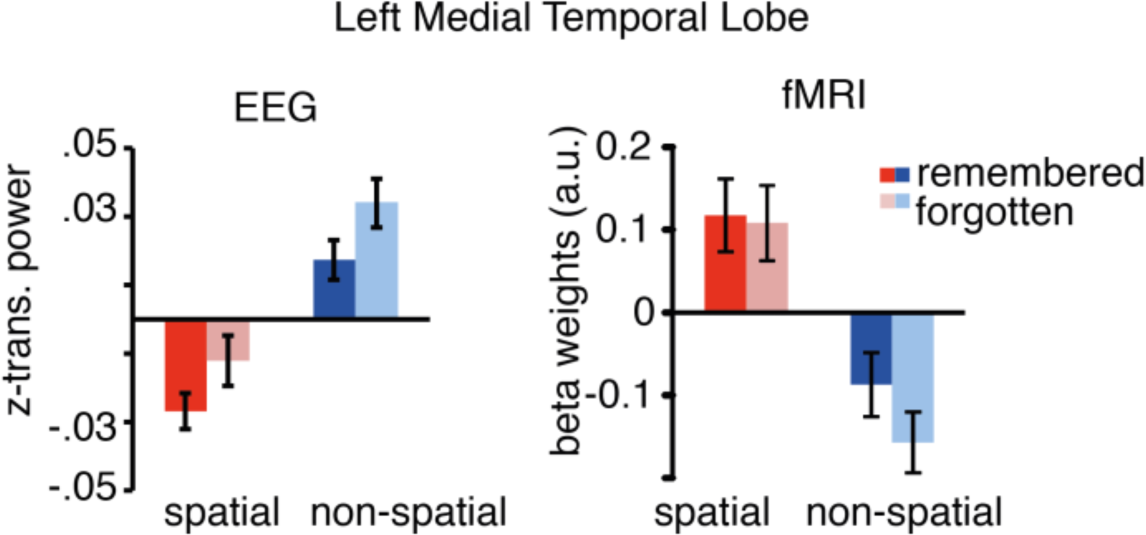
Mean EEG power and fMRI beta weights for the left MTL ROI. Power and beta weights were averaged for each condition. Theta power decreases show the reversed pattern of BOLD increases in left MTL regions Error bars show standard error of the mean.

## Discussion

Brain oscillations in the theta range and hemodynamic activity in the medial temporal lobe are considered to be core neural correlates of successful memory formation in electrophysiological and functional imaging studies, respectively. We here addressed the question of how theta power and MTL BOLD signals are functionally related. To this end we measured EEG and fMRI during two associative encoding strategies: the spatial Method of Loci, and the non-spatial Pegword Method. In line with previous work the spatial mnemonic indeed showed stronger increases in MTL BOLD signal relative to the non-spatial condition. Strikingly, these MTL BOLD signal increases were paralleled by MTL theta power decreases. Successful encoding during both mnemonics was predicted by equally pronounced left MTL BOLD signal increases, which were paralleled by theta power decreases in left lateral and medial temporal regions. Together, these findings suggest a negative relationship between theta power, memory formation, spatial processing, and MTL BOLD signals.

The Method of Loci recruits almost all cognitive processes that have been linked to MTL functions (i.e. scene construction, spatial processing, imagery, memory encoding and retrieval, and building new associative links (Bird and Burgess, 2008), which makes it ideal to study MTL theta oscillations during memory. Theta power decreases were source localized in areas overlapping with fMRI findings (see Table 1). Peaks of source localized theta power decreases were found in the parahippocampal gyrus and in left lateral temporal areas. Analysis of theta power changes placed in the right and left MTL ROIs closely followed the results of fMRI (see Figures 4&5). Spatial processing in contrast to non-spatial processing led to bilateral MTL theta decreases and BOLD increases. Subsequent memory effects in theta power and BOLD activity were evident in the left MTL. These parallels are remarkable and show that fMRI BOLD and EEG power can be linked via source localization techniques (Singh, 2012).

FMRI effects during spatial encoding closely follow previously reported effects relating to the usage of mnemonic strategies (Maguire et al., 2003) and spatial processing in general. The spatial mnemonic exhibited increases in BOLD signal in bilateral MTL and retrosplenial cortex, regions typically involved in imagining scenes and retrieving familiar landmarks (Burgess et al., 2002; Epstein, 2008). The Method of Loci task requires constructing spatial scenes to link to-be-learned items to familiar landmarks. Constructing such complex visual scenes crucially involves MTL structures (Addis et al., 2007; Hassabis and Maguire, 2007) with the retrosplenial cortex as a buffer of such imagined constructions (Bird and Burgess, 2008; Vann et al., 2009). The greater involvement of these brain structures in spatial processing, which additionally serve a central function in human memory (Hassabis and Maguire, 2007), might partly be responsible for the efficiency of the Method of Loci (Roediger, 1980).

Interestingly, subsequent memory effects in both mnemonics did not significantly differ depending on the spatial or non-spatial nature of the mnemonic, neither in EEG nor in fMRI. Memory formation during both tasks was related to activity in left MTL, consistent with the notion that the human MTL is involved in building item-cue associations (Ranganath, 2010; Buzsaki and Moser, 2013), regardless of the nature of these cues (i.e. spatial or non-spatial). Successful formation of associations in imagined navigational space and in semantic space thus seems to rely on similar mechanisms. This pattern of similar encoding effects for both encoding strategies supports the idea that humans use similar MTL mechanisms in spatial processing and in episodic memory encoding (see (Buzsaki and Moser, 2013; Ekstrom, 2014; Buffalo, 2015)).

Our findings with regards to posterior alpha/beta power increases matching BOLD signal decreases is well in line with several studies showing a negative relationship between low frequency power (<30 Hz) and BOLD signal (Mukamel et al., 2005; Hanslmayr et al., 2011; Scheeringa et al., 2011; Hermes et al., 2014; Zumer et al., 2014). These findings are compatible with several studies showing occipital alpha power increases to predict long term memory formation during internal processing of the memory material (Khader et al., 2010; Meeuwissen et al., 2011). Arguably, during the formation of these internal scenes the occipital cortex might be actively inhibited to prevent task-interfering visual input (Klimesch et al., 2007; Jensen and Mazaheri, 2010). Such an inhibition of posterior processing regions would be reflected by decreases in BOLD activity accompanied by alpha/beta power increases, which ultimately benefits memory.

Studies on oscillatory correlates of memory often implicitly assume a positive relationship between theta power increases and reported MTL BOLD increases (Summerfield and Mangels, 2005; Osipova et al., 2006; Staudigl and Hanslmayr, 2013; Backus et al., 2016), which is notable given that several intracranial EEG studies show of theta power decreases in the MTL during memory formation (Sederberg et al., 2007; Burke et al., 2013; Long et al., 2014; Greenberg et al., 2015). In line with these studies, and studies combining fMRI and electrophysiological measures (Mukamel et al., 2005; Niessing et al., 2005; Conner et al., 2011; Khursheed et al., 2011; Magri et al., 2012), the present results suggest that the negative relationship between low frequency power and BOLD also extends to theta oscillations in the MTL during memory encoding (Lisman and Jensen, 2013). The resemblance of low frequency power decreases during memory encoding in iEEG to positive SMEs in fMRI has been noted before (Burke et al., 2013) albeit no study before has shown this overlap in scalp EEG. Recording EEG and fMRI in the same paradigm, the present results therefore support a functional link between theta power decreases and MTL BOLD increases, showing that these neural processes not only occur in the same regions, but also co-vary with the same task conditions.

A central question that remains is how can we physiologically interpret theta power decreases paralleled by increases in BOLD signal? Traditionally, theta power increases have been hypothesized to reflect hippocampal-cortical feedback loops and increased neural cortico-hippocampal communication (Klimesch, 1996; Nyhus and Curran, 2010; Backus et al., 2016). Whether power increases are a suitable indicator of enhanced long-range neural communication however is questionable. For instance, intracranial recordings during memory formation reported decreases in low frequency theta power alongside increases in long-range phase synchrony (Burke et al., 2013). A similar relationship of local power decreases and global synchrony increases has been shown in other experiments (Popov et al., 2013; Weisz et al., 2014). Furthermore, theta oscillations which are dominant in the MTL and retrosplenial cortex during rest and movement (Ekstrom et al., 2009; Foster and Parvizi, 2012; Watrous et al., 2013b), might actually desynchronize during active tasks (Halgren et al., 1978; Mitchell et al., 2009), in order to flexibly form fine grained cortical networks connecting cortical regions to specific MTL sub-regions (Watrous et al., 2013a; Maris et al., 2016). Since the EEG records a spatially smoothed attenuated sum of LFPs (Buzsaki et al., 2012) fine grained long range synchronizations of specific individual cortical-MTL networks might actually appear as power decreases in scalp EEG and iEEG recordings.

A direct mapping of EEG power effects onto BOLD signals is complicated. Both modalities correlate with LFPs (Logothetis et al., 2001; Buzsaki et al., 2012) but do not necessarily reflect the same neural processes (Ekstrom, 2010). Adding to the differences of physiological correlates between EEG and fMRI, both modalities differ substantially in their temporal and spatial resolution. Considering these constraints, our EEG localization and fMRI results still exhibited a relatively good fit despite the low spatial resolution of EEG (Fuchs et al., 2002) and the low temporal resolution of fMRI. A limitation of the present study is that an analysis of simultaneously recorded theta oscillations during fMRI was not feasible due to strong artifacts induced by the MR scanning environment (Fellner et al., 2016). Therefore EEG and fMRI were measured separately in independent subject samples, preventing calculation of EEG-fMRI correlations. Nevertheless, the same paradigm was used in both datasets and the same behavioral pattern of results was found across the two datasets. Theta power changes and MTL BOLD changes were evident in the same task contrasts and were overlapping spatially. Therefore we can assume that, on average, the same cognitive processes were driving the EEG and fMRI effects in the two datasets (Mukamel et al., 2005; Singh, 2012).

## Conclusion

In summary, MTL and theta effects were more pronounced during the Method of Loci mnemonic, which indicates that navigating cognitive space might be a particularly efficient encoding strategy by maximally driving the neural processes related to spatial processing and episodic memory. The presented results show that decreases in theta oscillatory power in the MTL - similar to other cortical low frequency oscillations (i.e. alpha/beta) – co-occur with neural activity as reflected in the BOLD signal. MTL activity in memory tasks therefore seem to map onto decreases in EEG theta power and not, as often suggested, to increases in theta power.

## Acknowledgements

The research presented in this work was supported by a grant from the Deutsche Forschungsgemeinschaft (Project HA 5622/1-1) awarded to Simon Hanslmayr. Conflict of Interest: The authors declare no competing financial interests.

## References

Addis DR, Wong AT, Schacter DL (2007) Remembering the past and imagining the future: common and distinct neural substrates during event construction and elaboration. Neuropsychologia 45:1363–1377.

Backus AR, Schoffelen JM, Szebenyi S, Hanslmayr S, Doeller CF (2016) Hippocampal-Prefrontal Theta Oscillations Support Memory Integration. Curr Biol 26:450–457.

Bird CM, Burgess N (2008) The hippocampus and memory: insights from spatial processing. Nat Rev Neurosci 9:182–194.

Buffalo EA (2015) Bridging the gap between spatial and mnemonic views of the hippocampal formation. Hippocampus 25:713–718.

Burgess N, Maguire EA, O’Keefe J (2002) The human hippocampus and spatial and episodic memory. Neuron 35:625–641.

Burke JF, Zaghloul KA, Jacobs J, Williams RB, Sperling MR, Sharan AD, Kahana MJ (2013) Synchronous and asynchronous theta and gamma activity during episodic memory formation. J Neurosci 33:292–304.

Buzsaki G, Moser EI (2013) Memory, navigation and theta rhythm in the hippocampal-entorhinal system. Nat Neurosci 16:130–138.

Buzsaki G, Anastassiou CA, Koch C (2012) The origin of extracellular fields and currents–EEG, ECoG, LFP and spikes. Nat Rev Neurosci 13:407–420.

Coltheart M (2007) The MRC psycholinguistic database. The Quarterly Journal of Experimental Psychology Section A 33:497–505.

Conner CR, Ellmore TM, Pieters TA, DiSano MA, Tandon N (2011) Variability of the relationship between electrophysiology and BOLD-fMRI across cortical regions in humans. J Neurosci 31:12855–12865.

Ekstrom A (2010) How and when the fMRI BOLD signal relates to underlying neural activity: the danger in dissociation. Brain Res Rev 62:233–244.

Ekstrom A, Suthana N, Millett D, Fried I, Bookheimer S (2009) Correlation between BOLD fMRI and theta-band local field potentials in the human hippocampal area. J Neurophysiol 101:2668–2678.

Ekstrom AD (2014) Cognitive neuroscience: navigating human verbal memory. Curr Biol 24:R167–168.

Ekstrom AD, Caplan JB, Ho E, Shattuck K, Fried I, Kahana MJ (2005) Human hippocampal theta activity during virtual navigation. Hippocampus 15:881–889.

Epstein RA (2008) Parahippocampal and retrosplenial contributions to human spatial navigation. Trends Cogn Sci 12:388–396.

Fellner MC, Volberg G, Mullinger KJ, Goldhacker M, Wimber M, Greenlee MW, Hanslmayr S (2016) Spurious correlations in simultaneous EEG-fMRI driven by in-scanner movement. Neuroimage 133:354–366.

Foster BL, Parvizi J (2012) Resting oscillations and cross-frequency coupling in the human posteromedial cortex. Neuroimage 60:384–391.

Fuchs M, Kastner J, Wagner M, Hawes S, Ebersole JS (2002) A standardized boundary element method volume conductor model. Clin Neurophysiol 113:702–712.

Greenberg JA, Burke JF, Haque R, Kahana MJ, Zaghloul KA (2015) Decreases in theta and increases in high frequency activity underlie associative memory encoding. Neuroimage 114:257–263.

Halgren E, Babb TL, Crandall PH (1978) Human hippocampal formation EEG desynchronizes during attentiveness and movement. Electroencephalogr Clin Neurophysiol 44:778–781.

Hanslmayr S, Staudigl T (2014) How brain oscillations form memories–a processing based perspective on oscillatory subsequent memory effects. Neuroimage 85 Pt 2:648–655.

Hanslmayr S, Volberg G, Wimber M, Dalal SS, Greenlee MW (2013) Prestimulus oscillatory phase at 7 Hz gates cortical information flow and visual perception. Curr Biol 23:2273–2278.

Hanslmayr S, Volberg G, Wimber M, Raabe M, Greenlee MW, Bauml KH (2011) The relationship between brain oscillations and BOLD signal during memory formation: a combined EEG-fMRI study. J Neurosci 31:15674–15680.

Hassabis D, Maguire EA (2007) Deconstructing episodic memory with construction. Trends Cogn Sci 11:299–306.

Hermes D, Miller KJ, Vansteensel MJ, Edwards E, Ferrier CH, Bleichner MG, van Rijen PC, Aarnoutse EJ, Ramsey NF (2014) Cortical theta wanes for language. Neuroimage 85 Pt 2:738–748.

Jensen O, Mazaheri A (2010) Shaping functional architecture by oscillatory alpha activity: gating by inhibition. Front Hum Neurosci 4:186.

Kaplan R, Doeller CF, Barnes GR, Litvak V, Duzel E, Bandettini PA, Burgess N (2012) Movement-related theta rhythm in humans: coordinating self-directed hippocampal learning. PLoS Biol 10:e1001267.

Khader PH, Jost K, Ranganath C, Rosler F (2010) Theta and alpha oscillations during working-memory maintenance predict successful long-term memory encoding. Neurosci Lett 468:339–343.

Khursheed F, Tandon N, Tertel K, Pieters TA, Disano MA, Ellmore TM (2011) Frequency-specific electrocorticographic correlates of working memory delay period fMRI activity. Neuroimage 56:1773–1782.

Kim H (2011) Neural activity that predicts subsequent memory and forgetting: a meta-analysis of 74 fMRI studies. Neuroimage 54:2446–2461.

Klimesch W (1996) Memory processes, brain oscillations and EEG synchronization. Int J Psychophysiol 24:61–100.

Klimesch W, Sauseng P, Hanslmayr S (2007) EEG alpha oscillations: the inhibition-timing hypothesis. Brain Res Rev 53:63–88.

Klimesch W, Doppelmayr M, Russegger H, Pachinger T (1996) Theta band power in the human scalp EEG and the encoding of new information. Neuroreport 7:1235–1240.

Lisman JE, Jensen O (2013) The theta-gamma neural code. Neuron 77:1002–1016.

Logothetis NK, Pauls J, Augath M, Trinath T, Oeltermann A (2001) Neurophysiological investigation of the basis of the fMRI signal. Nature 412:150–157.

Long NM, Burke JF, Kahana MJ (2014) Subsequent memory effect in intracranial and scalp EEG. Neuroimage 84:488–494.

Magri C, Schridde U, Murayama Y, Panzeri S, Logothetis NK (2012) The amplitude and timing of the BOLD signal reflects the relationship between local field potential power at different frequencies. J Neurosci 32:1395–1407.

Maguire EA, Valentine ER, Wilding JM, Kapur N (2003) Routes to remembering: the brains behind superior memory. Nat Neurosci 6:90–95.

Maris E, Oostenveld R (2007) Nonparametric statistical testing of EEG-and MEG-data. J Neurosci Methods 164:177–190.

Maris E, Fries P, van Ede F (2016) Diverse Phase Relations among Neuronal Rhythms and Their Potential Function. Trends Neurosci 39:86–99.

Meeuwissen EB, Takashima A, Fernandez G, Jensen O (2011) Increase in posterior alpha activity during rehearsal predicts successful long-term memory formation of word sequences. Hum Brain Mapp 32:2045–2053.

Mitchell JF, Sundberg KA, Reynolds JH (2009) Spatial attention decorrelates intrinsic activity fluctuations in macaque area V4. Neuron 63:879–888.

Mukamel R, Gelbard H, Arieli A, Hasson U, Fried I, Malach R (2005) Coupling between neuronal firing, field potentials, and FMRI in human auditory cortex. Science 309:951–954.

Niessing J, Ebisch B, Schmidt KE, Niessing M, Singer W, Galuske RA (2005) Hemodynamic signals correlate tightly with synchronized gamma oscillations. Science 309:948–951.

Nyhus E, Curran T (2010) Functional role of gamma and theta oscillations in episodic memory. Neurosci Biobehav Rev 34:1023–1035.

Oostenveld R, Fries P, Maris E, Schoffelen JM (2011) FieldTrip: Open source software for advanced analysis of MEG, EEG, and invasive electrophysiological data. Comput Intell Neurosci 2011:156869.

Osipova D, Takashima A, Oostenveld R, Fernandez G, Maris E, Jensen O (2006) Theta and gamma oscillations predict encoding and retrieval of declarative memory. J Neurosci 26:7523–7531.

Popov T, Miller GA, Rockstroh B, Weisz N (2013) Modulation of alpha power and functional connectivity during facial affect recognition. J Neurosci 33:6018–6026.

Ranganath C (2010) A unified framework for the functional organization of the medial temporal lobes and the phenomenology of episodic memory. Hippocampus 20:1263–1290.

Roediger HL (1980) The effectiveness of four mnemonics in ordering recall. Journal of Experimental Psychology: Human Learning and Memory 6.

Scheeringa R, Fries P, Petersson KM, Oostenveld R, Grothe I, Norris DG, Hagoort P, Bastiaansen MC (2011) Neuronal dynamics underlying high-and low-frequency EEG oscillations contribute independently to the human BOLD signal. Neuron 69:572–583.

Schott BH, Henson RN, Richardson-Klavehn A, Becker C, Thoma V, Heinze HJ, Duzel E (2005) Redefining implicit and explicit memory: the functional neuroanatomy of priming, remembering, and control of retrieval. Proc Natl Acad Sci U S A 102:1257–1262.

Sederberg PB, Schulze-Bonhage A, Madsen JR, Bromfield EB, McCarthy DC, Brandt A, Tully MS, Kahana MJ (2007) Hippocampal and neocortical gamma oscillations predict memory formation in humans. Cereb Cortex 17:1190–1196.

Singh KD (2012) Which “neural activity” do you mean? fMRI, MEG, oscillations and neurotransmitters. Neuroimage 62:1121–1130.

Staresina BP, Davachi L (2009) Mind the gap: binding experiences across space and time in the human hippocampus. Neuron 63:267–276.

Staudigl T, Hanslmayr S (2013) Theta oscillations at encoding mediate the context-dependent nature of human episodic memory. Curr Biol 23:1101–1106.

Summerfield C, Mangels JA (2005) Coherent theta-band EEG activity predicts item-context binding during encoding. Neuroimage 24:692–703.

Uncapher MR, Otten LJ, Rugg MD (2006) Episodic encoding is more than the sum of its parts: an fMRI investigation of multifeatural contextual encoding. Neuron 52:547–556.

Van Veen BD, van Drongelen W, Yuchtman M, Suzuki A (1997) Localization of brain electrical activity via linearly constrained minimum variance spatial filtering. IEEE Trans Biomed Eng 44:867–880.

Vanderwolf CH (1969) Hippocampal electrical activity and voluntary movement in the rat. Electroencephalogr Clin Neurophysiol 26:407–418.

Vann SD, Aggleton JP, Maguire EA (2009) What does the retrosplenial cortex do? Nat Rev Neurosci 10:792–802.

Watrous AJ, Tandon N, Conner CR, Pieters T, Ekstrom AD (2013a) Frequency-specific network connectivity increases underlie accurate spatiotemporal memory retrieval. Nat Neurosci 16:349–356.

Watrous AJ, Lee DJ, Izadi A, Gurkoff GG, Shahlaie K, Ekstrom AD (2013b) A comparative study of human and rat hippocampal low-frequency oscillations during spatial navigation. Hippocampus 23:656–661.

Weisz N, Wuhle A, Monittola G, Demarchi G, Frey J, Popov T, Braun C (2014) Prestimulus oscillatory power and connectivity patterns predispose conscious somatosensory perception. Proc Natl Acad Sci U S A 111:E417–425.

Zumer JM, Scheeringa R, Schoffelen JM, Norris DG, Jensen O (2014) Occipital alpha activity during stimulus processing gates the information flow to object-selective cortex. PLoS Biol 12:e1001965.

